# Phloem anatomy restricts root system architecture development: theoretical clues from *in silico* experiments

**DOI:** 10.1101/2022.10.04.510862

**Authors:** Xiao-Ran Zhou, Andrea Schnepf, Jan Vanderborght, Daniel Leitner, Harry Vereecken, Guillaume Lobet

## Abstract

Plant growth and development involve the integration of numerous processes, influenced by both endogenous and exogenous factors. At any given time during a plant’s life cycle, the plant architecture is a readout of this continuous integration. However, untangling the individual factors and processes involved in the plant development and quantifying their influence on the plant developmental process is experimentally challenging.

Here we used a combination of computational plant models to help understand experimental findings about how local phloem anatomical features influence the root system architecture. In particular, we simulated the mutual interplay between the root system architecture development and the carbohydrate distribution to provide a plausible mechanistic explanation for several experimental results.

Our *in silico* study highlighted the strong influence of local phloem hydraulics on the root growth rates, growth duration and final length. The model result showed that a higher phloem resistivity leads to shorter roots due to the reduced flow of carbon within the root system. This effect was due to local properties of individual roots, and not linked to any of the pleiotropic effects at the root system level.

Our results open the door to a better representation of growth processes in plant computational models.

## Introduction

Plants develop complex architectures, both above ground (the shoot) and below ground (the roots) (Drouet and Pagès, 2003). An optimal architecture ensures that the different plant organs are ideally positioned in the environment to capture resources needed for their growth (Lobet et al., 2014b). Belowground, an efficient root system architecture ensures that individual roots are able to take up water and nutrients from the soil and transport them to the shoot (De Bauw et al., 2020). Since the availability of soil resources is highly heterogeneous, both in space and time, plants need to constantly adapt and develop their root system architecture (Morandage et al., 2021). Understanding how this development is controlled and regulated by the plant is an important open question.

The development of the root system fundamentally relies on three simple processes: root growth, self-re-orientation within the soil matrix (tropism) and production of next order (generation) of roots (Leitner et al., 2010). These new branches, or laterals, develop further following the same dynamical mechanisms. However, despite being individually simple, the assembly of these processes at the plant level quickly becomes complex as growth, tropism and branching are all influenced by exogenous and endogenous factors (Sharp et al., 2004; Ahmed et al., 2022). Exogenously, for instance, local soil water gradients can have a large influence on all the three processes (growth, tropism and branching). Endogenously, the amount of carbohydrates reaching individual root tips can restrict their growth. The amount of carbon reaching each growing tip is then directly linked to the growth potential and development of the whole root system.

Unlike leaves, roots are non-photosynthetic organs. The maintenance and growth at each location of a root system relies on the carbon that is transported from the leaves. This long-distance transportation from leaves to root takes place within the phloem vasculature, which is continuous throughout the whole plant. According to the pressure flow model of phloem transport, several key factors influence carbohydrates transport within the plant:

1. the loading rate of carbohydrates into the phloem sieve-tubes in source organs (e.g. leaves) (de Vries et al., 2021).
2. the unloading rate of carbohydrates out of the phloem sieve-tubes in sink organs (e.g. roots) (Uys et al., 2021).
3. the connections between the different plant organs (Delory et al., 2016).
4. the distribution of resistances of individual phloem vessels throughout the plant (Knoblauch et al., 2016).

How carbon transport controls growth is therefore a complex, multi-scale problem. Locally, the root phloem resistivity is influenced by its anatomy (i.e. the radius and number of the sieve-tubes as shown in Fig. S1); holistically, the length of a root changes its total resistance and influences the amount of available carbon at its tip (Fig. 1). The growth of a plant is therefore influenced by its architecture and anatomy, and defines both at the same time.

**Figure 1:**
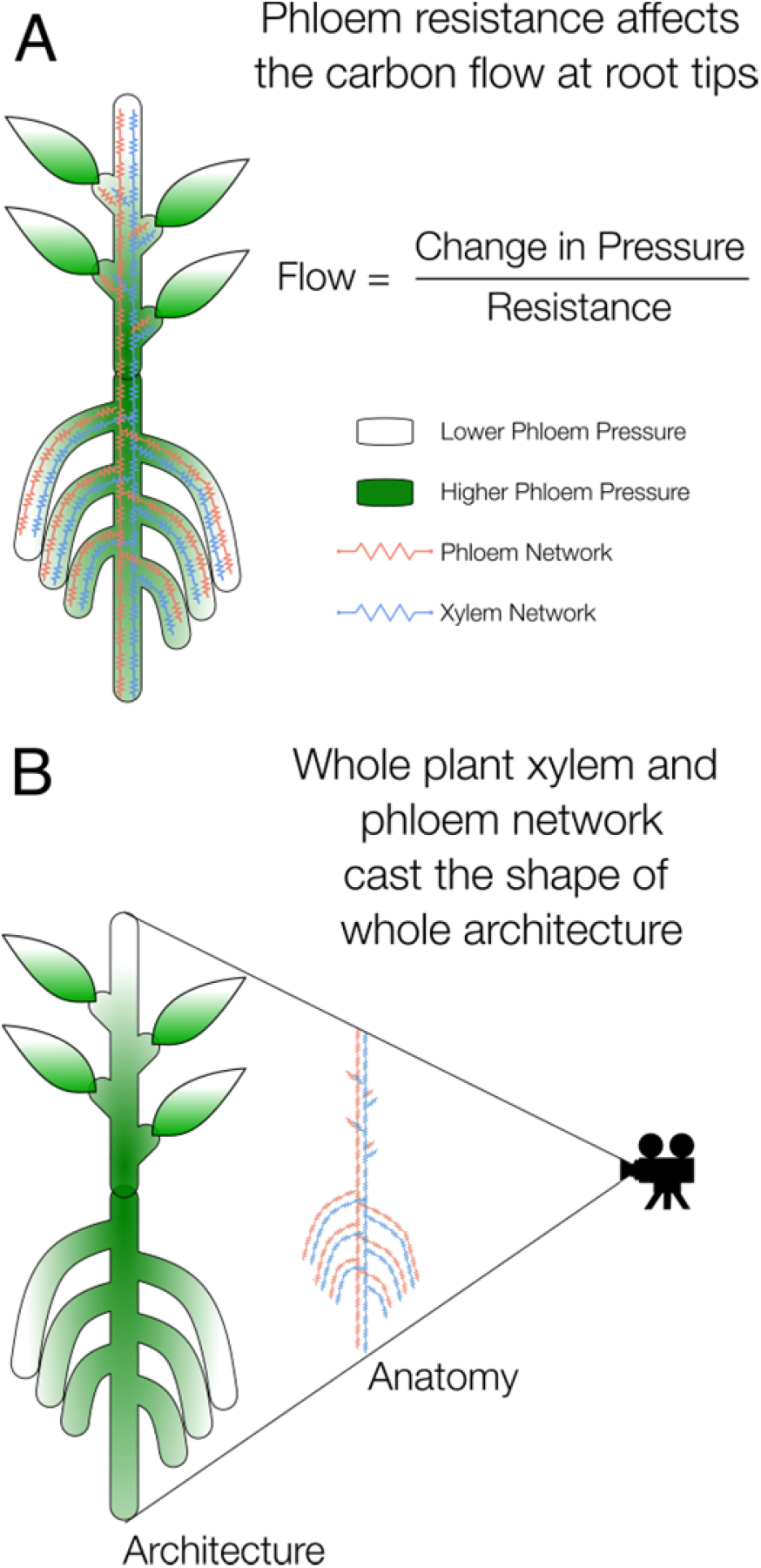
Plant vasculature casts the shape of architecture.**A**: The carbon flow reaching the end of the organ part is calculated through pressure and resistance. Thus, the phloem resistance affects the carbon flow at root tips. **B**: The limit of carbon content reaching the growing ends will also limit the growth. Thus, the axial resistivity of phloem, which is affected by anatomy, will also affect architecture. The anatomy can be used to cast the shape of plant architecture.

This creates a complex feedback loop between the phloem flow and the development of the root system. From an experimental point of view, disentangling the different processes (e.g. how resistivity of sieve-tubes affects root growth rate and final length) to understand their relative contribution is a complex task (Bidel et al., 2000; Dilkes et al., 2004).

Here we present an *in silico* analysis of such a complex system and further use this platform to explain experimental observations. To do so, we use a computational model of plant growth and development, CPlantBox (Zhou et al., 2020), coupled with a solver of the Pressure-Flow model of carbohydrates movement within the plant structure, PiafMunch (Lacointe and Minchin, 2008; Lacointe and Minchin, 2019). Using this model coupling, we show how carbohydrates transport and root system development are influencing each other and how they are influenced by local root phloem properties. In particular, we show how local phloem hydraulic properties can shape the root growth and development, and therefore modify the root system architecture as a whole.

## Results

### Overview of the analysis pipeline

In this study, we wanted to observe and quantify the theoretical effect of (1) sieve-tube resistivity on the local carbon flow within the root system and (2) the effect of the subsequent carbon (delivered by carbon flow) allocation on the root system development. To achieve this goal, we used a combination of state-of-the-art computational plant models.

First, we used the model CPlantBox (Zhou et al., 2020) to represent the plant structure. CPlantBox was designed to create a growing 3D plant structure (root and shoot), from a default user-defined parameter set. Second, we used a mechanistic model of phloem and xylem flow, PiafMunch (Lacointe and Minchin, 2019), to solve the distribution of carbohydrates within the plant. PiafMunch is able to predict the distribution of these carbohydrates within the whole plant structure, from a given plant structure, xylem and phloem hydraulic conductivities, and carbohydrate production rate,. Third, we created a dynamic feedback loop function between both models, such that the distribution of carbohydrates (in PiafMunch), at each time step, influences the formation of the plant architecture at the next time step (in CPlantBox). This allowed us to quantify the mutual effect between the distribution of carbohydrates and the establishment of the plant architecture.

The effect of the root phloem anatomy on the carbon distribution and root development was achieved via the modification of the phloem hydraulic conductivity (Zwieniecki et al., 2002; Jensen et al., 2012; Knoblauch et al., 2016; Holbrook and Knoblauch, 2018). Different phloem hydraulic resistance in both the primary root (study 1) and the lateral roots (study 2 and 3) lead to a different final root length. We also quantified the induced changes on the root system growth and development.

Details about the different parts of our analysis pipeline are given in the Material and Methods section.

### Phloem resistivities constrain the final length of primary and lateral roots

In the different simulations, we changed the phloem resistivity of the primary root or the phloem resistivities of the lateral roots. We observed a direct influence of such change on the final length of the affected roots (Fig. 2). As a general rule, when the phloem resistivity of a root increases, its final length decreases. This was observed for the primary roots (Fig. 2.A-B) and for the lateral roots (Fig 2.C-F).

**Figure 2:**
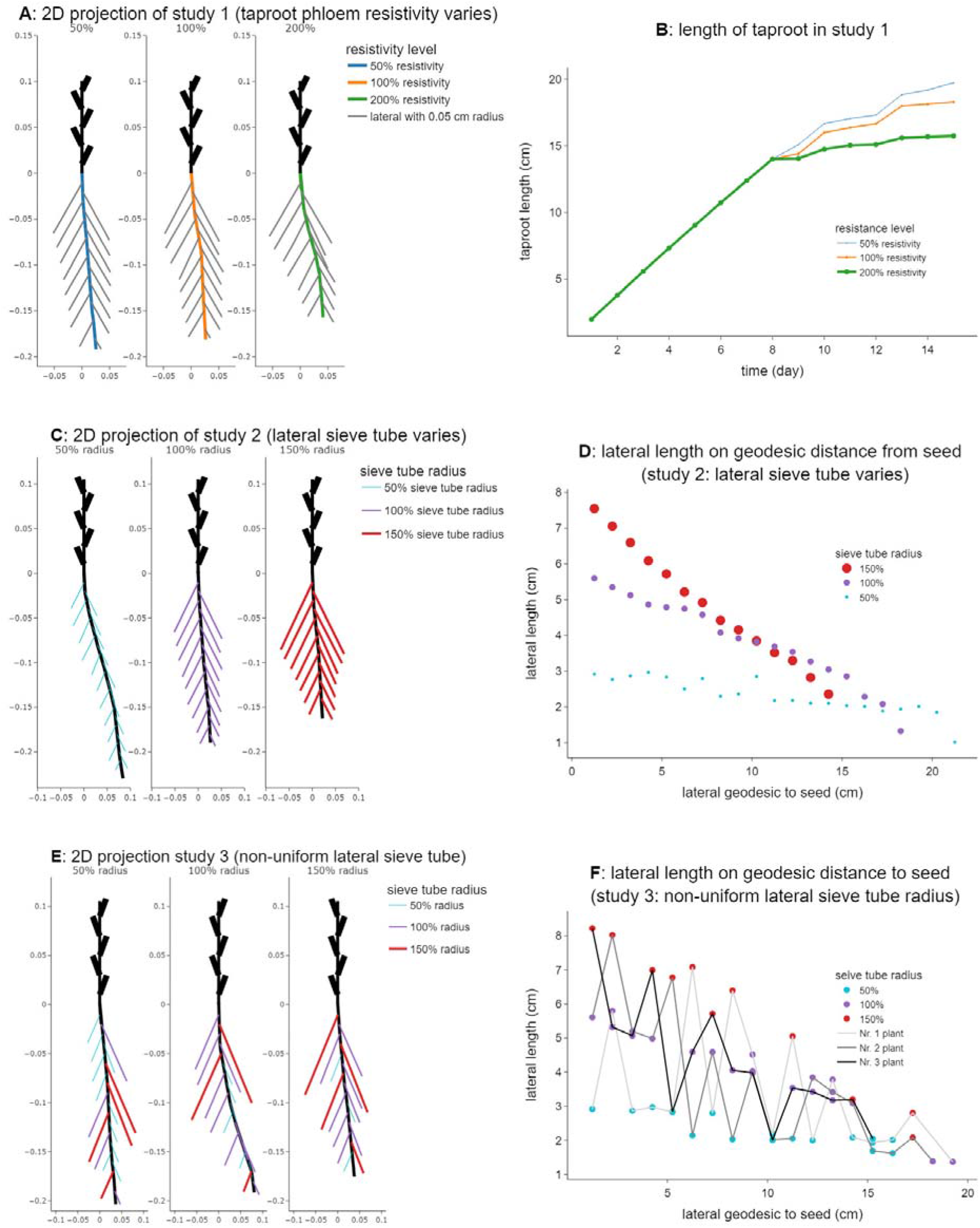
Growth patterns of study 1, 2 and 3 in 2-dimensional projections and length charts.**A**: 2D projection of study 1 (tap root sieve-tube varies); **B**: High resistant roots (thick green) grow less compared to low resistant roots (thin blue). **C**: 2D projection of study 2 (lateral sieve-tube varies). **D**: Different geodesic distance from seed to lateral base correlated to gradience of final lateral length. **E**: 2D projection study 3 (non-uniform laterals with different sieve-tube radius). **F**: Patterns of lateral length keep similar to study 2, even when laterals grow randomly.

In Fig. 2 E, F, we can also observe that the decrease in final root length is a consequence of a local change in root phloem resistivity and not a pleiotropic effect at the root system level. Indeed, when mixing roots with different phloem resistivities within a single root system (Fig. 2 E), we can see that the final length is influenced by these resistivities alone, and not the resistivities of the neighboring roots. Indeed, if we compare figures 2D and 2F, we can see that lateral roots with the same tube radius have similar behaviors, irrespective of their neighbors of position on the primary root.

### Phloem resistivity constrains the growth duration of primary and lateral roots

In addition to the effect of the phloem (single sieve-tube) resistivity on the final length of the roots, it seems to have a strong effect on the growth duration of individual roots (Fig. 3). In the model, the actual growth rate is indeed modulated by the amount of carbohydrates available at the root tip. If that amount is not enough to sustain the potential growth rate, as defined in the original parameter set, the growth rate is adjusted.

**Figure 3:**
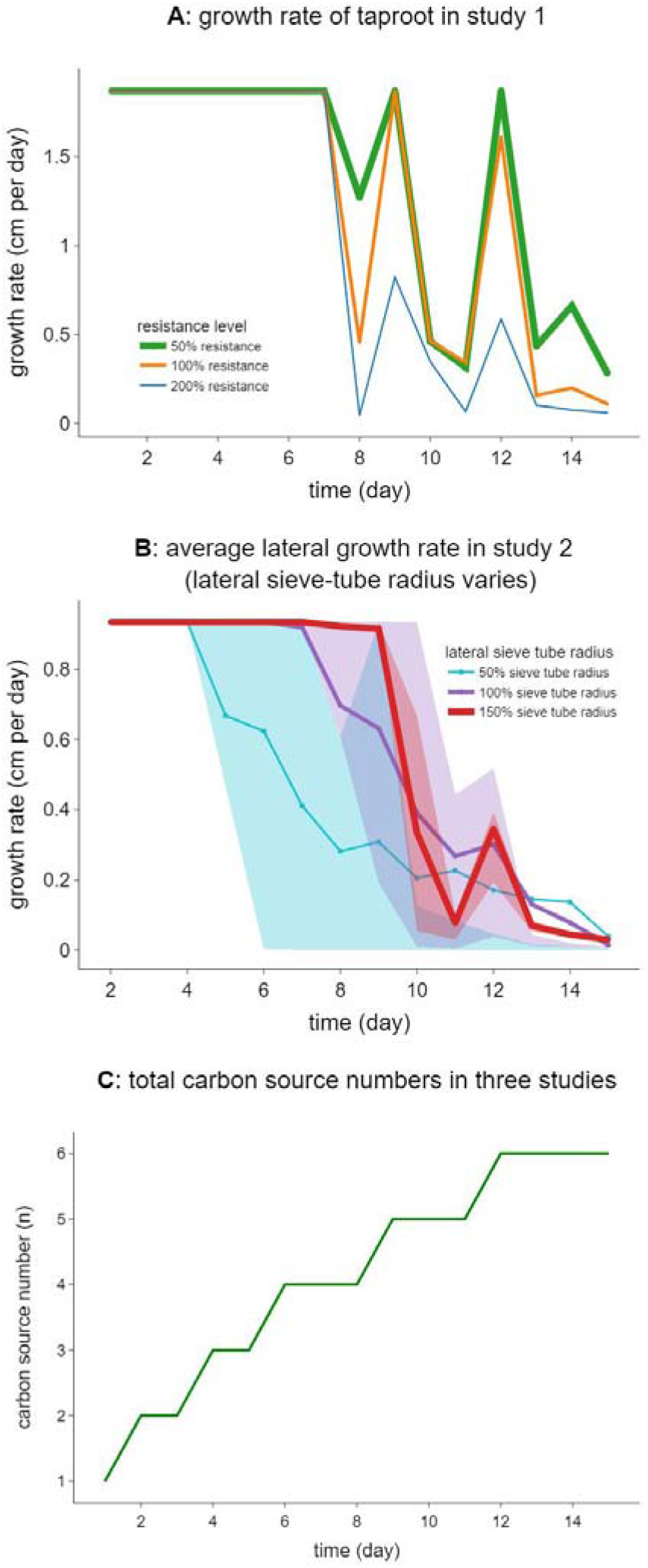
Root growth rate is controlled by both the phloem resistivity and carbon source input, while the final growth is only limited by the phloem resistivity.**A**: The increase of tap root growth rate on day 12 is caused by a newly growing leaf, which leads to more carbon loading into the phloems; **B**: Average (colored lines) and quantile (25% to 75% in color area) of the laterals’ growth rates in study 2; **C**: On day 6, 9 and day 12, we can see some tips’s growth rate increased comparing to previous day. Compared to the growth on day 6, the growth on day 12 is much less. This indicates that the additional carbon source (a newly growing leaf) is no longer the deciding factor, when the phloem resistance is limiting the growth.

Again, for both the primary and the lateral roots, we observed that the changes in phloem resistivity impose strong constraints on duration of growth. For the tap roots, the growth is identical for the first 7 days, independently of the phloem resistivity (Fig. 3 A). After the 7th day, the root with the largest resistivity decreases its growth, then stops after the 15th day completely. The same can be observed after 7 days for the roots with 100% and 50% single sieve-tube resistivity, respectively.

For the lateral roots, the picture is less clear, as many roots are influenced at the same time. However, the same dynamic can be observed as for the primary root. Indeed, the duration of the growth period seems to be directly linked to the hydraulic resistivity of the phloem (Fig. 3B). As the phloem resistivity of individual roots increases, the growth period of these roots shortens.

### Local phloem resistivities influence carbon allocation between the root and the shoot

The observed changes in growth can be linked to changes in the carbohydrate flow within the root systems. In the different simulations, the total carbon production was equal across the three studies (same loading rate of carbohydrates in the phloem at the shoot level) (Fig. 4 A). However, the allocation between the root and the shoot is different in the different studies (Fig. 4 B, C). We can indeed observe that in the study with the largest primary root resistance, the root system carbon content is lower, leading to a smaller root-to-shoot ratio.

**Figure 4:**
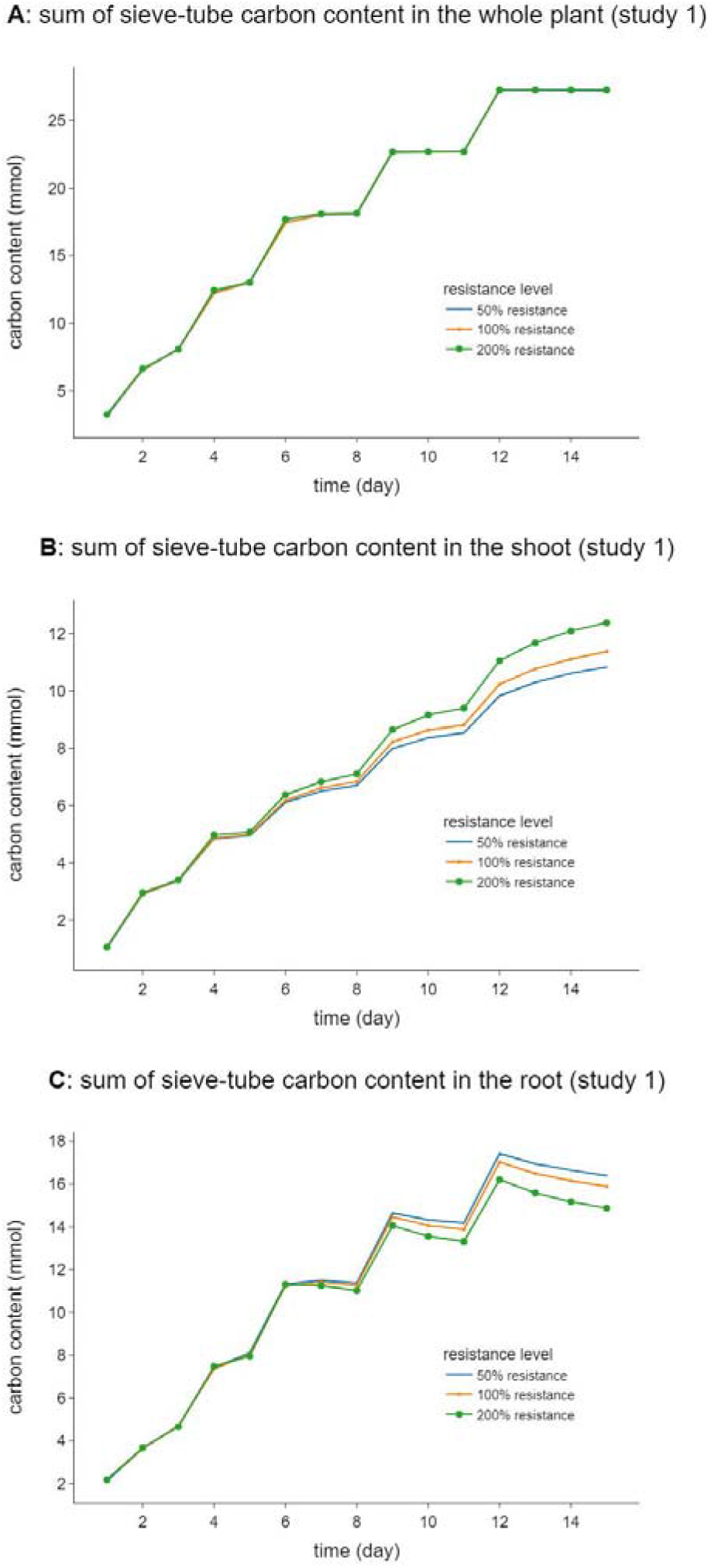
Comparison among the sums of sieve-tube content of whole plants, shoots and roots in study 1.**A**: After the 8th day, the sum of all sieve-tube solute carbon contents in the whole plants are the same between plants, although which have different taproot phloem resistivities. The slight difference mainly showed up during the first eight days, might be due to the excessive carbon supply. There is slightly more carbon in the phloem of higher resistant plants, because more carbon needs to be kept in the phloem to generate a higher pressure to push the carbon down to the root. **B**: In the shoot, leaves are growing, which in turn increase the sieve-tube carbon content. **C**: In the root, the sums of sieve-tube carbon content in the three plants are similar during the first 6 days, because the root sizes are similar. Later on, the root with larger resistivity grows less, in turn the plant with high phloem resistance has smaller root size and contains less carbon content. It is very interesting to see that the carbon content in sieve-tubes reached equilibrium by distributing carbon content between the shoots (sources) and roots (sinks).

The allocation patterns are not predefined in the model, but the output of the mechanistic model which describes the carbon flow within the system. The changes in allocation are therefore a direct consequence of the altered phloem resistivities.

## Discussion

### Phloem resistance creates a sink limitation to carbohydrate flow

In our simulations, we could observe a strong limitation to the carbon flow in the system due to changes in the conductivities along the carbon pathways (and not directly due to changes in local carbon sink strength). As the carbon loading at the leaf level was set at a constant value in all simulations, the total amount of carbohydrates injected into the system was constant for each leaf (and therefore increasing during the simulation as new leaves were produced Fig. 3C). However, the distribution (allocation) of the carbohydrates within the plant was strongly influenced by total root phloem resistances. In our simulations, the phloem anatomy was therefore directly linked to a sink limitation of carbon flow within the plant. As the root phloem resistance along the path of carbohydrates becomes too high, the flow decreases to a stop, preventing the resources, which are needed for individual root growth, to reach the root apexes. It is worth noting that, in our simulations, the limitation is not at the sink itself (the potential demand for carbon was identical in all simulations), but on the path of carbon leading to that sink. If that path is limiting the flow, it becomes impossible to push carbohydrates along the roots, no matter how strong the local demand at the apexes is. This brings a new perspective to previous studies that have shown a strong effect of local root tip demand on the global architecture, but did not take the resistance along the phloem vasculature explicitly into account (Pagès et al., 2020).

From a breeding perspective, this indicates that the phloem resistance in the roots can be a strong restriction to root/shoot allocation manipulation. If we aim at creating larger root systems, either to access more resources (Lobet et al., 2012; Lynch, 2013; Lobet et al., 2014a) or to store mode carbon within the soil (Bidel et al., 2000; Dilkes et al., 2004), it is critical to take this sink limitation into account.

### Phloem anatomy as a driver of root architecture

Our simulation results indicate that, in theory, local phloem resistance can have a strong influence on the growth and development of single roots and therefore influence the shape and function of the root system as a whole. The effects observed in this study are limited to small plant architectures but are expected to become larger as the plant grows. Small effects observed at the beginning of the growing period will accumulate during the root system development, leading to very distinct developmental trajectories.

This opens up interesting perspectives in root research. It means that some specific phloem developmental traits, such as to locally increase or decrease its resistivity, has the potential to manipulate the growth and development of specific root types and to shape the root system architecture as a whole.

From a root modeling perspective, our results suggest that final root length of specific roots could be seen as output of the model, rather than inputs, as it is the case with most current root models (Pagès et al., 2004; Pagès et al., 2014; Postma et al., 2017; Schnepf et al., 2018). However, the modeling approach described here is likely to be too demanding in terms of computing resources, making it impractical for large scale modeling studies. A tighter integration between the different models (here CPlantBox and PiafMunch) or the use of upscaling methods could help to incorporate these mechanistic processes into larger scale models.

### Phloem anatomical variance explains uneven development between roots with different diameters

Many studies have established links between root diameter and function. For instance, root diameter has been shown to be linked to specific root length and mycorrhizal colonization (Ma et al., 2018), foraging capacity (Colombi et al., 2017) or the root capacity to uptake water (Heymans et al., 2020). Root diameter has also been linked to root developmental properties such as growth rate (Pagès and Picon-Cochard, 2014), branching density (Pagès, 2016) or root system architecture as a whole (Pagès and Kervella, 2018). These links have been shown to hold true between species, but also within single plants. Recent studies have quantified the growth and development of several root types growing on the same primary roots (Passot et al., 2016; Passot et al., 2018; Muller et al., 2019). Recently, a clear connection between phloem diameter and growth was shown by (Tang et al., 2022). These different root types have been shown to have contrasting anatomy, growth rate and final length, leading to highly heterogeneous root system architectures, even within homogeneous conditions.

We were able to qualitatively reproduce various distinct developmental dynamics (Clerx et al., 2020; Tang et al., 2022) (differences in growth rate and final length) by changing only the root phloem resistivities. As phloem resistivity is, to some extent, linked to the root diameter (as the diameter increases the number of phloem poles increases as well), our simulation results could explain some of the differences observed between roots of different diameters (Passot et al., 2016; Passot et al., 2018; Muller et al., 2019).

## Conclusion

In this study, we have used a computational pipeline to explore the interplay between local phloem hydraulic properties and the global root system development. We have shown that a local decrease in the phloem conductivity can lead to shorter roots, with a slower growth rate and shorter growth duration. We have also shown that this change in architecture was not due to a pleiotropic effect at the plant level, but mainly a local effect of the phloem properties.

Our results are well in line with experimental results that have studied the link between root diameter, length, and growth. They provide a simple mechanistic explanation to the variety of root types observed experimentally and provide an alternative explanation to the effect of root tip sinks alone.

Finally, from a modeling point of view, our results provide a first example where the features of single roots within the root system (such as their growth rate and final root length) are not prescribed *a priori* but are an emergent property of the system. This opens the way to move plastic representation of the root system within computational models.

## Authors contributions

Writing—original draft: X.-R .Z., G.L.; writing—review and editing: J.V., H.V., G.L., X.-R .Z.; conceptualization: G.L., A.S., H.V., J.V., X.-R .Z.; software: X.-R .Z., D.L., A.S., G.L.; visualization: X.-R .Z., G.L., J.V.

## Code availability

The code used in this paper is available here [10.5281/zenodo.7115421] and here [https://github.com/Plant-Root-Soil-Interactions-Modelling/CPlantBox/tree/zhou2022]

## Acknowledgement

This research was supported by Agrosphäre, IBG-3 Forschungszentrum Jülich. We thank our colleagues from UMR PIAF who provided insight and expertise that greatly assisted the research, although they may not agree with all of the interpretations of this paper.

We thank André Lacointe for assistance with PiafMunch. We would also like to show our gratitude to the other members of ROSI group for sharing their pearls of wisdom with us during the course of this research, and we thank the anonymous reviewers (in advance).

## Material and methods

### Brief descriptions of the models CPlantBox and PiafMunch

CPlantBox is a generic computational plant model (Zhou et al., 2020) and it was designed to create a growing plant structure containing both the root and the shoot based on growth rules that describe organ elongation, branching, orientation, and senescence. CPlantBox can be connected to other models, in such a way that the simulated architecture can be used as a numerical grid to simulate diverse functions such as water flow (Morandage et al., 2021), exudation (Landl et al., 2021), nutrient uptake (De Bauw et al., 2020) or carbon flow (Zhou et al., 2020).

PiafMuch (Lacointe and Minchin, 2008; Lacointe and Minchin, 2019) is a solver of coupled carbon and water flow in phloem and xylem respectively, following Münch Theory (Münch, 1930). In both models, PiafMunch and CPlantBox, the topology of the plant is abstracted as a set of connected nodes in 3D space. On top of the topology, functional parameters such as axial resistance of xylem and phloem, maintenance, loading rate and unloading rate are assigned on each node or connection. From this information, PiafMunch simulates the advective transport of carbohydrates through the phloem within the whole structure and returns the amount of carbon (delivered by sieve-tube flow) available for unloading at any node. Water flow in both xylem and phloem follows the Hagen-Poiseuille’s law. The hydraulic water potentials drive both the xylem and phloem water flow, while only the phloem water flow with dissolved carbon is affected by the pressure generated by the osmotic gradients (Münch Hypothesis). There is water flow exchange between the xylems and phloems, which depends on the pressure and resistance between xylem and phloem, throughout the simulation. The carbon unloading is described as a first-order kinetic process in which the unloading rate is linearly dependent on the carbon concentration in the sieve-tubes at each root tip. Excess carbon (unloaded carbon not used for the growth of the root tip) is considered to be exuded.

Here, we connected both models, such that, at each time step, the plant architecture is computed by CPlantBox, sent to PiafMunch, that in turn sends back to CPlantBox the amount of carbon available for growth at each root tip. The development of the structure in CPlantBox at the next time step is therefore constrained by the available carbon. The dynamic feedback loop between both models is illustrated in Fig. 5 and explained below.

**Figure 5:**
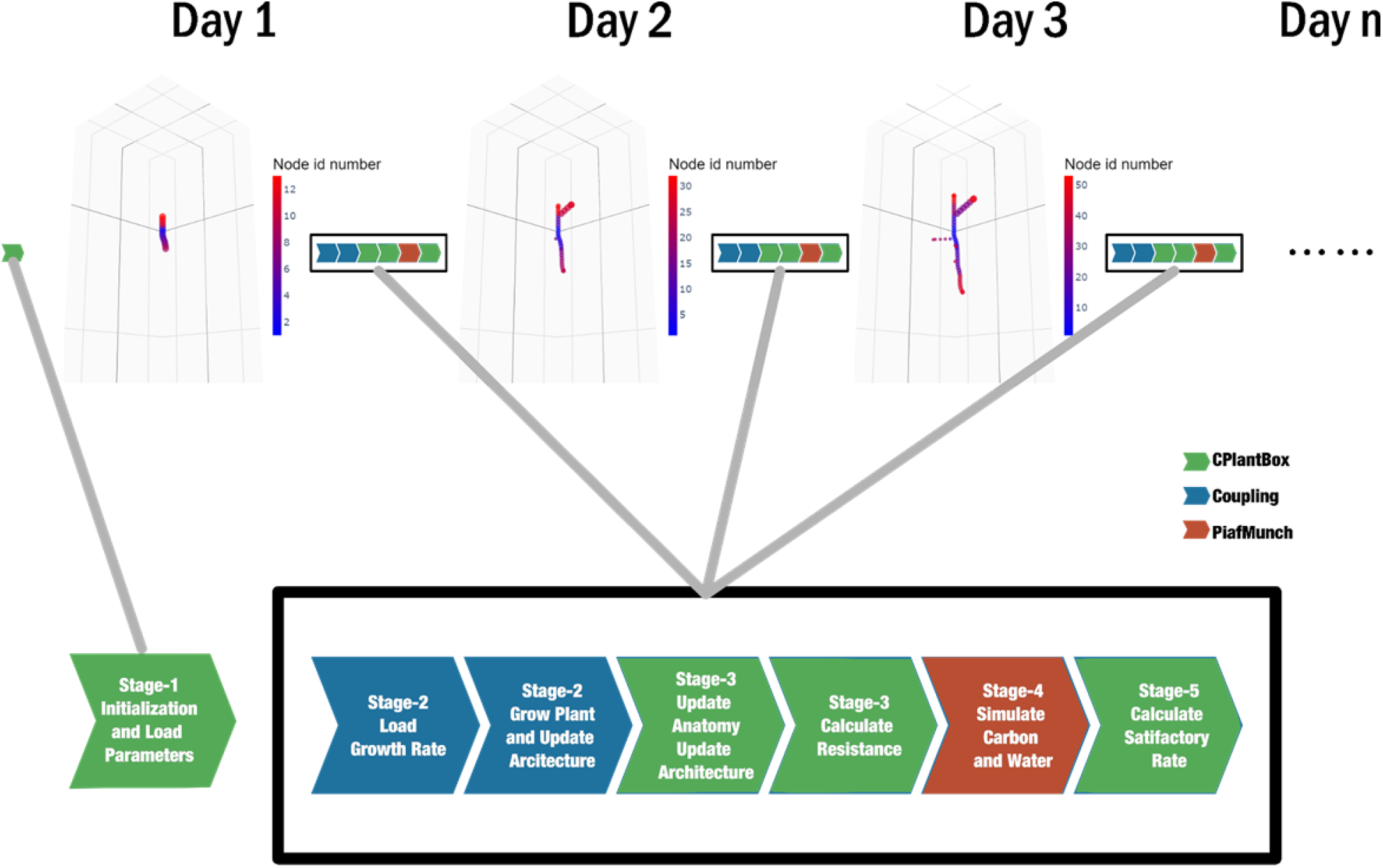
Feedback loop between CPlantBox and PiafMunch (detailed description in section dynamic feedback between structures and functions, code is open-sourced in https://github.com/Plant-Root-Soil-Interactions-Modelling/CPlantBox/blob/870e70d99276483173f520f03b7496a19c4b8978/tutorial/jupyter/zhou_2022/2022_publication_simulation_scripts.ipynb)

### Dynamic feedback between structures and functions

A Jupyter Notebook (Perkel, 2018) was used as a mid-ware to control both CPlantBox (run under Windows Subsystem for Linux) and PiafMunch (run under Windows), see Section Code Availability. The Jupyter Notebook was used to initialize parameters and run simulations.

#### Stage 1: Initialization of parameters for CPlantBox and PiafMunch

Most parameters of CPlantBox, such as organ radius, potential growth rate and potential topological structure of each plant are initialized and kept constant throughout the simulation, except for the root growth rate, which is either pre-defined in the previous studied (Zhou et al., 2020) or depends on the carbon delivered to the root tip (this study). Although it is possible to generate some stochasticity by adding deviation to the parameters, in this paper we did not use any stochasticity except the random emergence of laterals in study 3. Previously, the XML parameter file applied the same growth rate to all the laterals with the same sub-type (Zhou et al., 2020). Considering carbon availability, the growth rate may be different between each single growing root tip, resulting in different final lengths.

Similar to the initialization of CPlantBox, most parameters of PiafMunch are also kept constant throughout each study except the architecture and xylem resistances, which changes as shown in supplemental Fig. S2 (every 24 hours). Phloem resistivity and sieve-tube radius of specific root types, maximum loading rate, maximum unloading rate of each source or sink are kept constant in each study. Details about the estimation of the different parameters are given in Supplemental Material 1.

#### Stage 2: Generation of the plant structure

Unlike the initialization stage, which happens only once per study, the root architecture grows at every time-step. For simplicity, we assume that the shoot growth rate is not affected by carbon or water. However, the actual root growth rate is calculated for each root tip, depending on the potential growth rate (as defined in the parameter file, which is constant among each *sub-type*) and the available carbon (as defined by PiafMunch in the previous time step). The actual growth rate is therefore always equal or lower than the potential growth rate as it may be limited due to the local carbon availability.

#### Stage 3: Coupling between CPlantBox and PiafMunch

At stage three in each time step, the architecture simulated by CPlantBox is sent to PiafMunch, and carbon and water flow are computed for this architecture. The hydraulic properties of each root segment are computed based on their topology and geometry. While the root xylem radial and axial resistances are computed based on their local segment age (Doussan et al., 1998; Heymans et al., 2021), the sieve-tube axial resistances are assumed to be constant within root types and not age dependent. They are calculated according to the Poiseuille law for water flow in a cylindrical tube, i.e., they are inversely proportional to the fourth power of the sieve-tube radius (see Equation 1), see also supplemental Fig. S1).

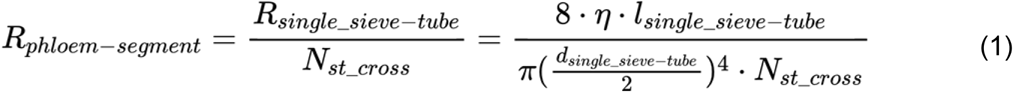

In Equation 1, *R*_*phloem-segment*_ (MPa h ml^-1^) is the resistance of a phloem segment, *R*_*single_sieve-tube*_ (MPa h ml^-1^) is the resistance of a single phloem sieve-tube, N_st_cross_ (dimensionless) is the number of sieve-tubes in a root cross section (dimensionless), η is the viscosity of the phloem sap, which is assumed to be constant as 1.7 (mPs · s), *l*_*single_sieve-tube*_ (cm) is the length of the root segment, *d*_*single_sieve-tube*_ (cm) is the diameter of a single sieve-tube.

The dry root mass of each segment is calculated by the following equation:

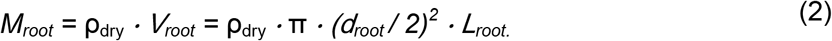

In Equation 2, V_root_ (cm^3^) is the volume of one root segment, d_root_ (cm) is the diameter of the root segment, L_root_ (cm) is the length of this root segment and ρ_dry_ (g cm^-3^) is the dry mass density of root (which is set to 0.1 (g cm^-3^) according to (Drouet and Pagès, 2003)). The total carbon of a segment used for maintenance is set to zero.

#### Stage 4: Simulating water and carbon flow

At stage 4, PiafMunch takes the architecture and anatomy information generated in stage 3, to simulate the carbon and water flow as explained in (Zhou et al., 2020). Although only carbon is studied in all the three studies, water is also simulated in PiafMunch. This is because xylem water flow is a prerequisite of the carbon flow, see (Minchin and Lacointe, 2017) for details. At the end of each time step, an output file containing the distribution of carbon within the plant structure is created and sent back to CPlantBox.

#### Stage 5: Limiting root growth rate with local carbon availability

In CPlantBox, the available carbon limits the potential growth rate of individual roots via a satisfactory rate coefficient. The carbon satisfactory rate is calculated as the ratio between the carbon needed for the potential growth rate (as defined in the CPlantBox parameter set) and the local carbon availability, as defined by PiafMunch, at each time step. The actual growth rate is therefore calculated by multiplying the potential growth rate (as defined in the input parameter file) by the carbon satisfactory rate. Each individual root tip uses the corresponding actual growth rate, *g*_*actual*_ (cm day^-1^), in the next time step. The calculation is based on the following equation,

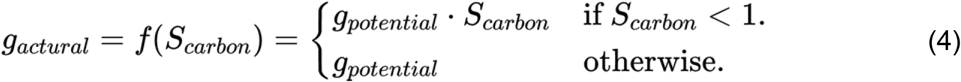

where *S*_*carbon*_ is the carbon satisfactory rate (a ratio); *g*_*potential*_ is the potential growth rate (cm day^-1^). The calculation of *S*_*carbon*_ (dimensionless cm cm^-1^) can be found in Supplemental Material 1

### Description of the study setups

We created three *in silico* studies to investigate the effect of local root anatomies on the formation of the root system architecture. In each study, three “treatments” were simulated. In all three treatments in each study, all the structural and functional parameters are kept the same, unless stated otherwise. The simulated plant is a simple plant, with a single taproot bearing first order lateral roots. The shoot growth and development is the same in all three study simulations and not influenced by the carbon distribution within the plant. A description of the three studies is given below and summarized in table 1.

**Table 1.** Comparison of the sensitivity studies

|  | Taproot | Lateral root |  |
| --- | --- | --- | --- |
|  | Single sieve-tube resistivity | Single sieve-tube resistivity | Mixed lateral roots |
| **Study 1** | **varied** | fixed | no |
| **Study 2** | fixed | **varied** | no |
| **Study 3** | fixed | **varied** | **yes** |

#### Study 1: Variations in tap root resistivity

In the first study, we quantified the effect of taproot phloem resistivity changes on the carbon flow toward the root system and the consequence on the root system development. To do so, we only changed the taproot phloem resistivity between the plants of this study.

#### Study 2: Variations in lateral sieve-tube radius

In the second study, we quantified the effect of the heterogeneous resistivity in lateral roots (induced by the heterogeneity of sieve-tube radii) on the carbon flow toward the root system and the consequence on the root system development. In this study, we therefore change the initial lateral root sieve-tube diameter (which is constant throughout the study) for the different treatments.

#### Study 3: Non-uniform lateral sieve-tube radius

In the third study, we quantified the effect of the lateral root sieve-tube radius heterogeneity on the carbon flow toward the root system and the consequence on the root system development. In contrast with study 2, in this study three types of lateral roots with different sieve-tube radii are grown in random order on the taproot in each study.

